# Taxonomy of *Pseudomonas* spp determines interactions with *Bacillus subtilis*

**DOI:** 10.1101/2023.07.18.549276

**Authors:** Mark Lyng, Birta Þórisdóttir, Sigrún H. Sveinsdóttir, Morten L. Hansen, Gergely Maróti, Lars Jelsbak, Ákos T. Kovács

## Abstract

Bacilli and pseudomonads are among the most well-studied microorganisms commonly found in soil, and frequently co-isolated. Despite this, no systematic approach has been employed to assess the pairwise compatibility of members from these genera. Here, we screened 720 fluorescent soil isolates for their effects on *Bacillus subtilis* pellicle formation in two types of media and found a predictor for interaction outcome in *Pseudomonas* taxonomy. Interactions were context-dependent and both medium composition and culture conditions strongly influenced interactions. Negative interactions were associated with *Pseudomonas capeferrum, Pseudomonas entomophila* and *Pseudomonas protegens*, and 2,4-diacetylphloroglucinol was confirmed as a strong (but not exclusive) inhibitor of *B. subtilis*. Non-inhibiting strains were closely related to *Pseudomonas trivialis* and *Pseudomonas lini*, but in this case, cocultures with increased *B. subtilis* pellicle formation were spatially segregated. Our study is the first to propose an overall negative outcome from pairwise interactions between *B. subtilis* and fluorescent pseudomonads, hence cocultures comprising members from these groups are likely to require additional microorganisms for coexistence.

## Introduction

Plant growth-promoting rhizobacteria (PGPR) possess great potential to replace traditional fertilisers and pesticides as a more sustainable alternative. In particular, isolates from *Bacillus* and *Pseudomonas* genera have been studied extensively due to their abilities to inhibit plant pathogens, induce plant systemic resistance, increase the growth rate of plants, and alleviate environmental stress (*Bacillus* reviewed in ^1^ and *Pseudomonas* reviewed in ^2^). Several studies combined isolates from the two genera and observed a synergistic increase in a trait of interest, be it plant growth or protection ^3–7^. However, few studies have investigated the cause of synergy, hence little information is available on the pairwise compatibility of *Bacillus* and *Pseudomonas*, and whether environmental isolates engage in antagonism or symbiosis.

*Bacillus subtilis* is one of the most well-studied model organisms in biology, serving as a prototypical example of biofilm formation and plant root colonisation. Several strains have demonstrated antagonism towards other microorganisms, especially plant pathogenic fungi ^8^. Such interactions are mainly mediated by secreted bioactive secondary metabolites, of which *B. subtilis* produces a diverse arsenal. Of note is plipastatin that shows antifungal properties ^9,10^, while surfactin and bacillaene display more broad antimicrobial properties ^11,12^. Positive interactions between *Bacillus* and *Pseudomonas* have also been observed, such as intraspecies division of labour ^13^ and interspecies cross-feeding ^14^.

*Pseudomonas* is a genus of numerous species comprising at least 166 type strains that are found in soil, plant rhizospheres, marine habitats and animal hosts ^15–18^. The impact of *Pseudomonas* spp. on human society ranges from human and plant pathogenicity to bioremediation, biocontrol and biostimulation ^19– 22^. Interestingly, there is a fine line between pathogenic and beneficial *Pseudomonas* spp. that mainly depends on the arsenal of secondary metabolites produced by a given strain ^23,24^. Similarly, compatibility with *Bacillus* spp. also seems to rely mainly on a relatively small collection of secondary metabolites and defence mechanisms ^25^. Understanding the frequency of positive and negative interactions between *Bacillus* and diverse *Pseudomonas* spp., as well as the underlying mechanisms of such interactions would be of great interest to agricultural biotechnology relying on mixed consortia of species from the two genera.

Here, we cocultured 720 soil isolates selected for fluorescent *Pseudomonas* properties with the undomesticated type-strain *B. subtilis* DK1042 (hereafter DK1042) under floating biofilm (pellicle)-inducing conditions and used a spinning disc-based medium-throughput screen to quantify DK1042 biofilms. Using two types of media, we demonstrate that although positive interactions with superior pellicle abundance were rare in both types of media, interactions were highly medium-dependent. By taxonomically characterising isolates at the species level, we determined that DK1042 was least likely to be compatible with species closely related to type strains of *Pseudomonas capeferrum, Pseudomonas entomophila* and *Pseudomonas protegens*, and most likely to be compatible with *Pseudomonas lini* and *Pseudomonas trivialis*. We found that *Pseudomonas* antagonism towards DK1042 is mainly due to the presence of specific biosynthetic gene clusters (BGCs), and that 2,4-diactylphloroglucinol (DAPG) is a major (though not exclusive) inhibitor of DK1042 against both pellicles and colonies.

## Results

### Pseudomonads negatively affect *B. subtilis*

Relationships between *B. subtilis* and fluorescent pseudomonads range from antagonism to co-existence and, potentially, to synergistic growth, but no systematic investigation of interaction outcomes between the two has been performed. To address this, we performed pairwise coculture of DK1042 and each of 720 *Pseudomonas* soil isolates in 96-well plates and measured DK1042 biomass using an Opera High-Content screening spinning disc platform (Fig. 1a). We screened the library against DK1042 *amyE*::P_hyperspank_-*mKate2* constitutively expressing the fluorophore mKate2, and recorded the three-dimensional volume occupied by *Bacillus* as a proxy for cell abundance. Each isolate was cocultured three times with DK1042 both in rich Tryptic Soy Broth (TSB) medium and diluted biofilm-inducing lysogeny broth medium supplemented with glycerol and manganese (0.1× LBGM), and biovolume in coculture was compared with that in monocultures as log_2_(Fold Change) (Fig. 1b).

**Fig. 1.**
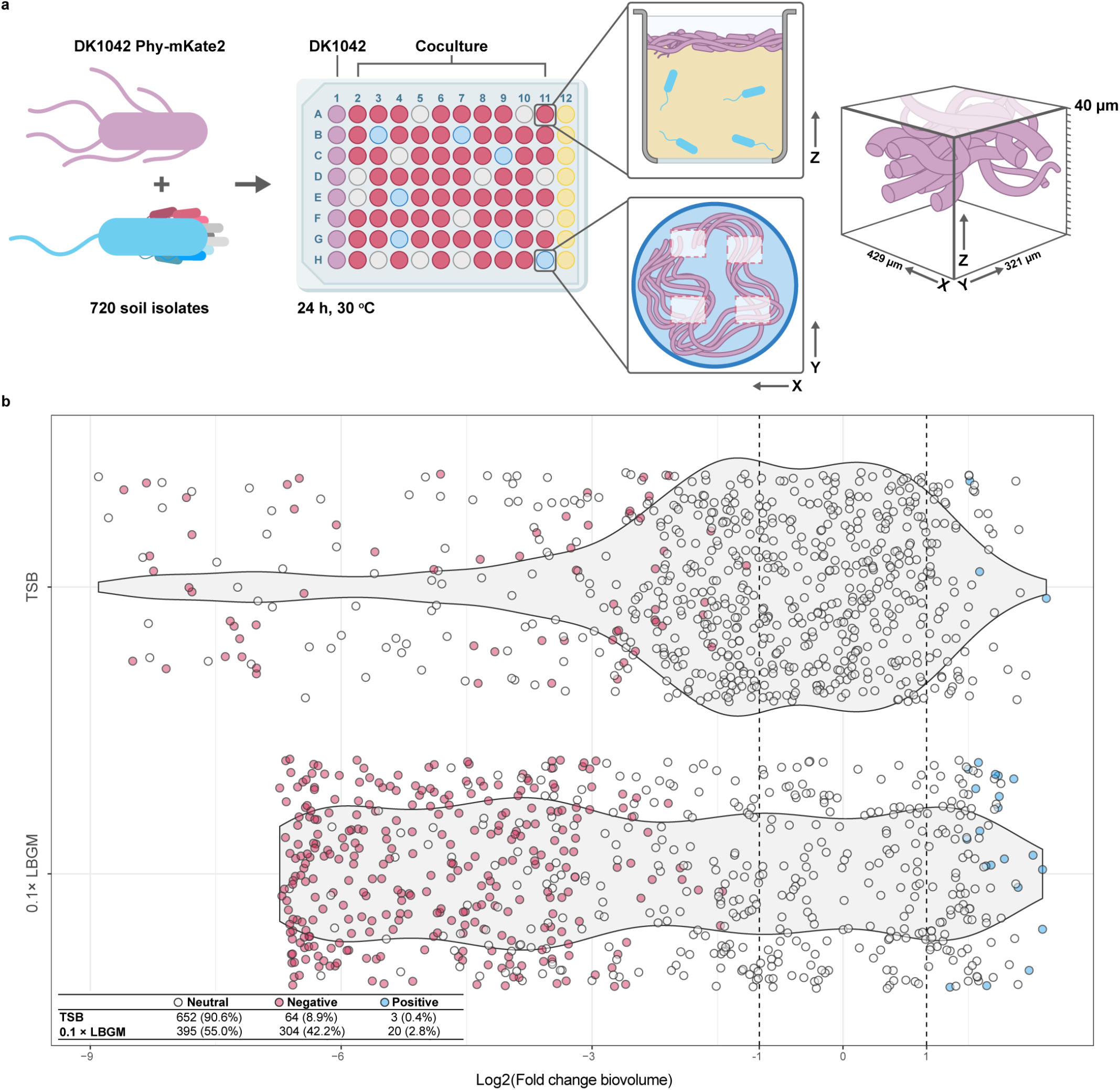
Pseudomonads negatively affect *B. subtilis*. **a:** DK1042 constitutively expressing mKate2 was mixed pairwise with 720 fluorescent soil isolates and cultured for 24 h at 30°C in a microplate format. Monocultures of DK1042 (purple wells) were used for comparison of pellicle formation, and wells with non-inoculated medium (yellow wells) were used to control for contamination. Each well was imaged in four positions with a Perkin Elmer Opera QEHS, acquiring images in the Z-direction every 2 µm to obtain a cube with height 40 µm. Media was removed prior to microscopy. **b:** DK1042 biovolumes were compared between co- and monoculture to yield log_2_(Fold Change). Points represent medians of three replicates. Neutral, negative and positive categories were assigned based on median and interquartile range.

The distribution of log_2_(FC) differed substantially between media types, suggesting that many neutral interactions in rich media become negative in diluted media, in line with stronger resource competition in nutrient-poor media among microorganisms ^26^. Examining the biovolume for each isolate (Fig. S2a) revealed that isolates from site P9 were more likely to antagonise DK1042 while isolates from the P8 site more often resulted in a high DK1042 biovolume. This also differed between media types, as cocultures in TSB had a higher degree of within-sample variance in biovolume measurements and therefore also in calculated log_2_(FC) values (Fig. S2b and S2c). We divided each isolate into three categories based on median log_2_(FC) and found that most interactions (regardless of medium) were neutral (Fig. 2a), while only a very small number of isolates had a positive effect on DK1042 (Fig. 1b). These results indicate that fluorescent pseudomonads very rarely stimulate *B. subtilis* cell abundance in TSB or 0.1× LBGM, and that inhibition of *B. subtilis* by *Pseudomonas* spp. is strongly influenced by medium composition.

**Fig. 2.**
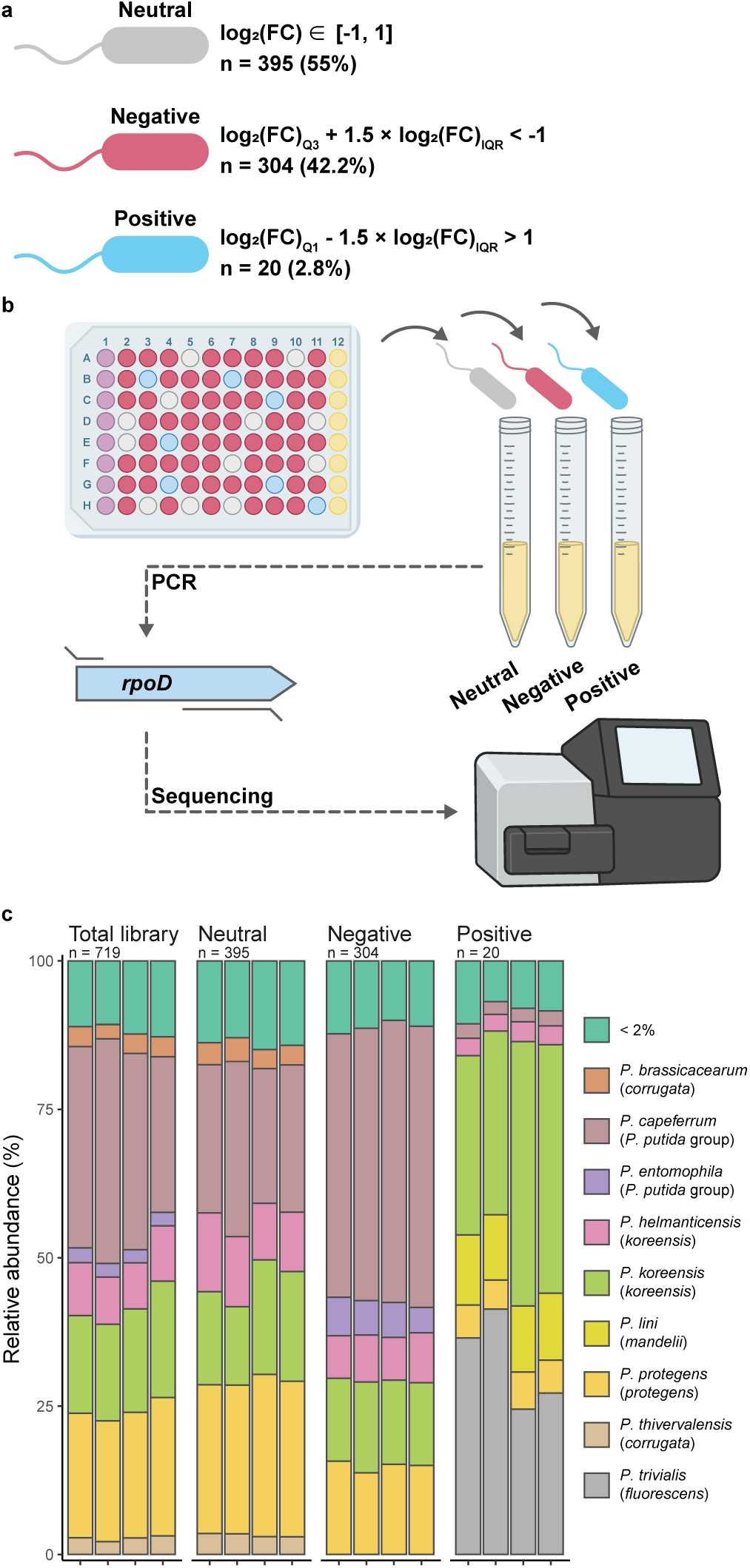
Screening categories enriched for species-specific taxa. **a:** Fluorescent isolates were assigned a category based on median and interquartile range, such that negatives resulted in DK1042 log_2_(FC) <-1 and positives in log_2_(FC) >1. n and percentages are from screening in 0.1× LBGM. **b:** Isolates were grown in precultures and pooled in equal cell numbers according to their categorisation to taxonomically characterise each category at the species level via *rpoD* amplicon sequencing. **c:** Relative abundance of *Pseudomonas* spp. in each category. Parentheses indicate groups or *P. fluorescens* subgroups (see Table 1 for details on taxonomy). Taxons comprising <2% of a pool were merged.

### Pseudomonas taxonomy predicts interaction outcome

To determine the taxonomic distribution of isolates, we sequenced amplicons of the *rpoD* gene from pools of isolates corresponding to their screening category (neutral, negative and positive; Fig. 2 and Dataset S1).

**Table 1:** Significantly differentially abundant species in screening categories compared with the total library.

| *fluorescens*<br>subgroup<sup>a</sup> | *Pseudomonas* sp. | Neutral<br>(log<sub>2</sub>FC) | Negative<br>(log<sub>2</sub>FC) | Positive<br>(log<sub>2</sub>FC) |
| --- | --- | --- | --- | --- |
| *asplenii* | *P. asplenii* | 1.363* | -0.235 | -0.153 |
| *corrugata* | *P. thivervalensis* | -0.028 | -0.090 | -0.921* |
|  | *P. brassicacearum* | -0.066 | -0.263 | -1.123* |
| *fluorescens* | *P. trivialis* | **-0.292** | **-0.381** | **2.900\*\*** |
|  | *P. cedrina* | -1.607* | 0.646 | -1.110 |
|  | *P. libanensis* | -0.084 | -0.038 | -1.124* |
|  | *P. lurida* | 0.153 | -1.097 | -1.196* |
| *gesaardii* | *P. brenneri* | -1.176 | 1.048 | -1.261* |
| *koreensis* | *P. helmanticensis* | 0.063 | 0.173 | -0.884* |
|  | *P. granadensis* | 0.341 | -1.736* | -1.134 |
|  | *P. moraviensis* | 0.031 | -0.267 | -1.244* |
| *mandelii* | *P. lini* | **-0.048** | **-0.425** | **2.623\*\*** |
|  | *P. migulae* | -0.510 | -0.348 | 1.829* |
|  | *P. frederiksbergensis* | -0.163 | -1.292 | 1.574** |
|  | *P. mandelii* | 0.088 | 1.007* | 0.396 |
| *protegens* | *P. protegens* | **-0.015** | **-0.090** | **-1.215\*\*** |
| *P. putida*<sup>b</sup> | *P. mosselii* | -0.935 | 1.213** | -0.830 |
|  | *P. soli* | -1.558* | 0.856 | -1.443* |
|  | *P. entomophila* | -2.064** | 1.138* | -2.294** |
|  | *P. capeferrum* | **-0.452** | **0.627** | **-2.493\*\*** |

<sup>a</sup> As described in <sup>15</sup>.
<sup>b</sup> Not a subgroup of *P. fluorescens*.
\* Differentially abundant from the total library with FDR-adjusted *p*-value <0.05. \*\* FDR-adjusted *p*-value <0.01.
**Bold** indicates predictions.

The species diversity of each category demonstrates how the categorisation based on *B. subtilis* pellicle formation selectively distributes specific phyla into separate categories (Permutational Analysis of Variance, *p* <0.001, Fig. S3). The total library of 720 isolates was mainly comprised of *P. capeferrum, Pseudomonas helmanticensis, Pseudomonas koreensis* and *P. protegens* (Fig. 2c and Table 1). A similar distribution occured in the neutral category, while negative and positive strains differed significantly in species abundance. The pool of negative isolates contained more strains associated with the *P. putida* group compared with the total library, but no single species was significantly depleted from this category. Differential abundance analysis revealed that the pool of positive isolates contained fewer isolates from the *Pseudomonas putida* group, and the *P. protegens, Pseudomonas corrugata* and *P. koreensis* subgroups, compared with the total library (Table 1). By contrast, *P. trivialis* and members of the *Pseudomonas mandelii* subgroup were enriched. As the pool of positive strains only contained 20 isolates, the statistical power in determining differential abundance was quite low. Therefore, caution should be applied when inferring from statistical tests in this group, hence we refer to isolates henceforth as “inhibiting” (negative) or “non-inhibiting” (neutral and positive).

To test the predictions resulting from screening, we obtained four *Pseudomonas* strains closely related to species that were implicated in different interaction patterns, and cocultured them with DK1042 in liquid and solidified 1× LBGM and 0.1× LBGM media (Fig. 3). As predicted, *P. capeferrum* and *P. protegens* both inhibited pellicle formation (even in 1× LBGM), while neither *P. lini* nor *Pseudomonas poae* (in place of *P. trivialis*) reduced pellicle abundance or winkle formation (as measured via pixel standard deviation; Fig. 3b). However, on solid media, no pseudomonad was able to reduce the colony size of DK1042 on 1× LBGM, but on diluted 0.1× LBGM *P. protegens* strongly antagonised DK1042

**Fig. 3.**
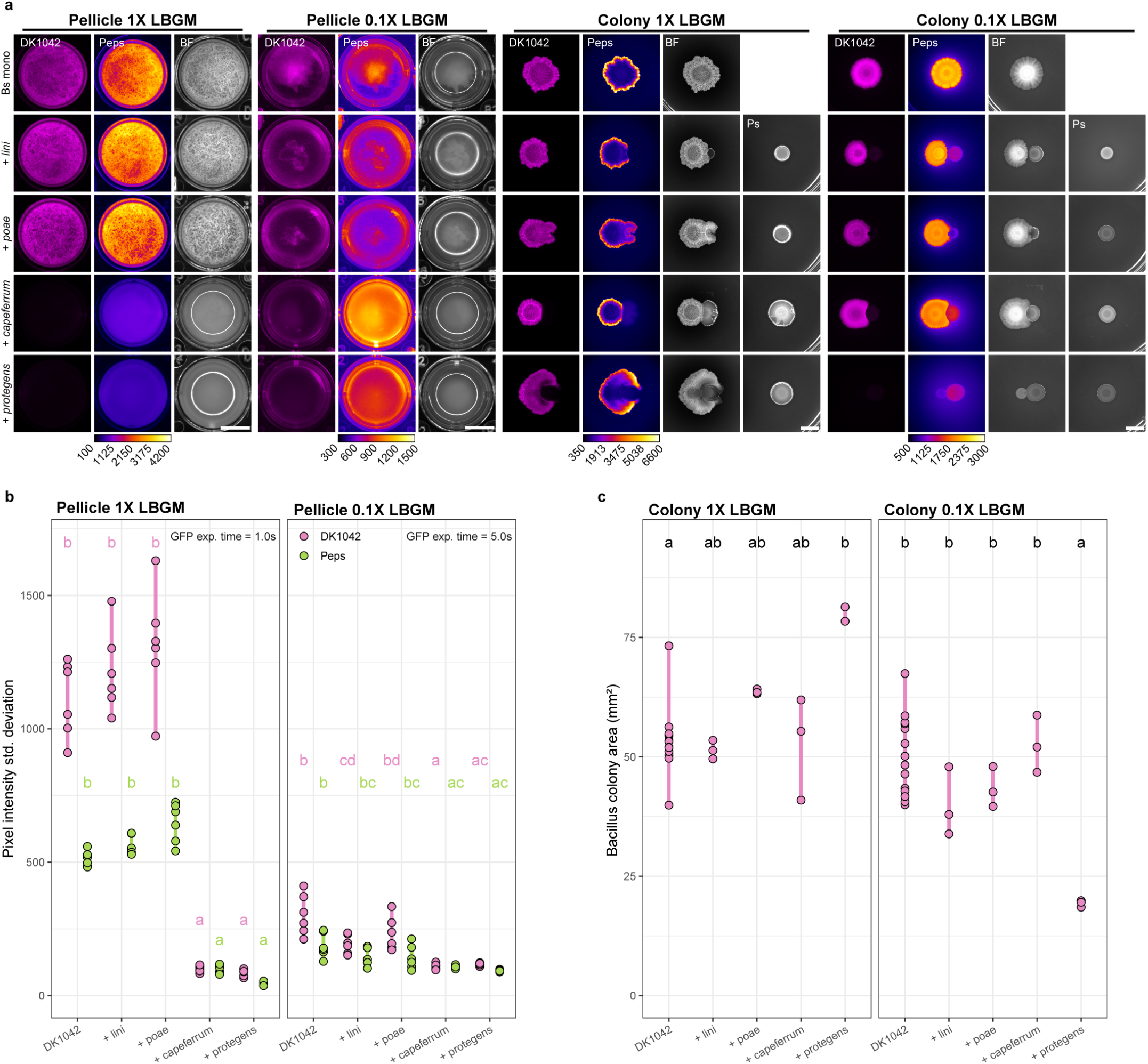
Screening prediction of inhibitors and non-inhibitors. **a:** Cocultures of DK1042 with four reference strains that were independent from the soil isolate library were used to assess the validity of interaction prediction resulting from screening *Pseudomonas* soil isolates. Cultures were grown at 30°C for 24 h (pellicles) or 72 h (colonies). Cultures were prepared with DK1042 reporting expression of the *epsA−O* operon using GFP. Scale bar = 5 mm. **b:** Pellicle wrinkle formation represented by the standard deviation in pixel intensity (i.e. high standard deviation equals stronger wrinkle formation). n = 6 independent experiments. **c:** Colony area of DK1042 spotted as monoculture or neighbouring the four *Pseudomonas* reference strains. n = 3 independent cocultures. Grouping letters are from ANOVA with Tukey-Kramer’s post-hoc test. Identical letters within each plot indicate a statistically significant grouping (*p* <0.05).

(Fig. 3c). Thus, *Pseudomonas* antagonism of DK1042 not only depends on medium constituents, but also on mode of growth.

### BGC abundance does not determine interaction outcome

*Pseudomonas* secondary metabolites are often implicated in *Bacillus*-*Pseudomonas* antagonism ^12,27,28^, therefore we sequenced the genomes of 13 candidate isolates and predicted BGCs and BGC subgroups using antiSMASH (Fig. 4). Interestingly, there was no significant difference in the abundance of encoded BGCs between inhibiting and non-inhibiting isolates (Fig. 4a). Therefore, we examined each genome for the presence of BGCs with known products (Fig. 4b). As expected, all isolates carried a version of pyoverdine, a fluorescent siderophore, but many non-inhibitory isolates were negative for most other biosynthetic gene clusters compared with inhibiting isolates. Isolates from *P. mandelii* and *Pseudomonas jesenii* subgroups collectively encoded only one BGC with a predicted product (rhizoxin, originally isolated from *Paraburkholderia rhizoxinica* ^29^). By contrast, colony-inhibiting isolates from *P. corrugata* and *P. protegens* subgroups both carried genes encoding 2,4-diacetylphloroglucinol(DAPG)-producing enzymes, while *P. protegens* additionally encoded enzymes for producing pyochelin, orfamide A, pyrrolnitrin, and pyoluteorin.

**Fig. 4.**
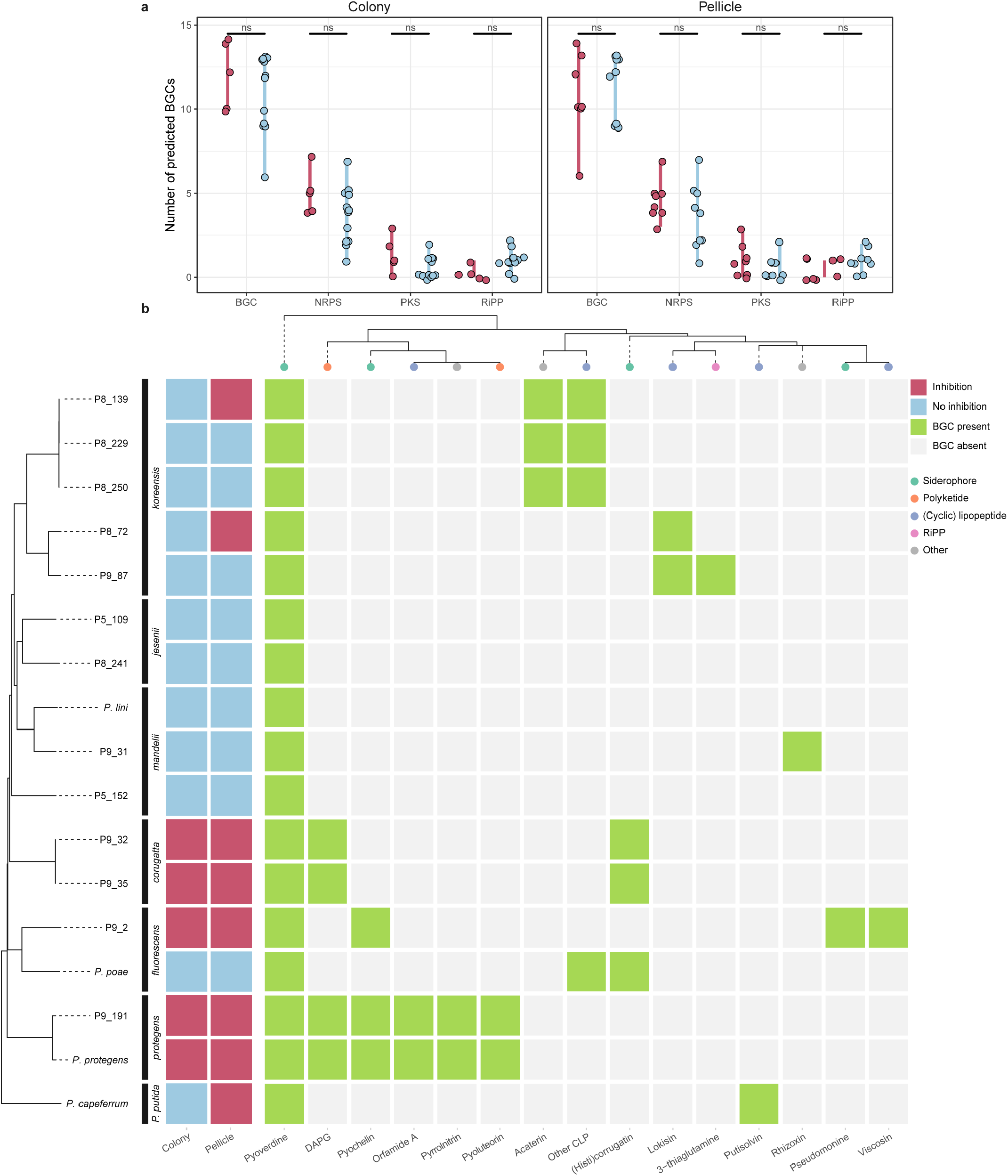
Specific BGCs (not abundance) predict interaction outcome. Thirteen isolates were whole-genome sequenced and compared with the four reference strains. **a:** AntiSmash predictions of biosynthetic gene clusters (BGCs) and BGC subgroups grouped by inhibition potential in colony or pellicle cocultures. Statistical tests were pairwise Student’s t-tests adjusted for multiple testing with the false discovery rate method. ns, not significant (adj. *p* >0.05). **b:** *Pseudomonas*-related natural product machinery encoded within each strain. Strain phylogeny is based on whole-genome sequence identity as described in TYGS ^76^.

### Non-inhibiting isolates are spatially segregated from DK1042

While the main biofilm mode of *B. subtilis* in liquid culture involves growth at the air-liquid interface ^30^, *Pseudomonas* spp. are usually observed colonising submerged surfaces in laboratory environments, although examples of *Pseudomonas* pellicle formation do exist ^31,32^. The presence of pellicles in cocultures with non-inhibiting isolates therefore suggests three potential scenarios: either non-inhibiting pseudomonads are outcompeted by *B. subtilis*, are growing with *B. subtilis* in the pellicle, or are spatially isolated from *B. subtilis* on the submerged surface of the microtiter plates. To determine which of these scenarios occurred with DK1042, we fluorescently labelled a non-inhibitor (P9_31), cocultured it statically with DK1042 in liquid 1× LBGM and 0.1× LBGM, and imaged the entire depth of the well (Fig. 5). In both media, P9_31 was spatially segregated to the submerged surface, even when DK1042 was mostly present in the pellicle. Interestingly, while cultivation in 1× LBGM resulted in a pellicle both with and without P9_31, cultivation in 0.1× LBGM required P9_31 for pellicle formation, even though the *Pseudomonas* isolate was present at the bottom of the well and not in the pellicle.

**Fig. 5.**
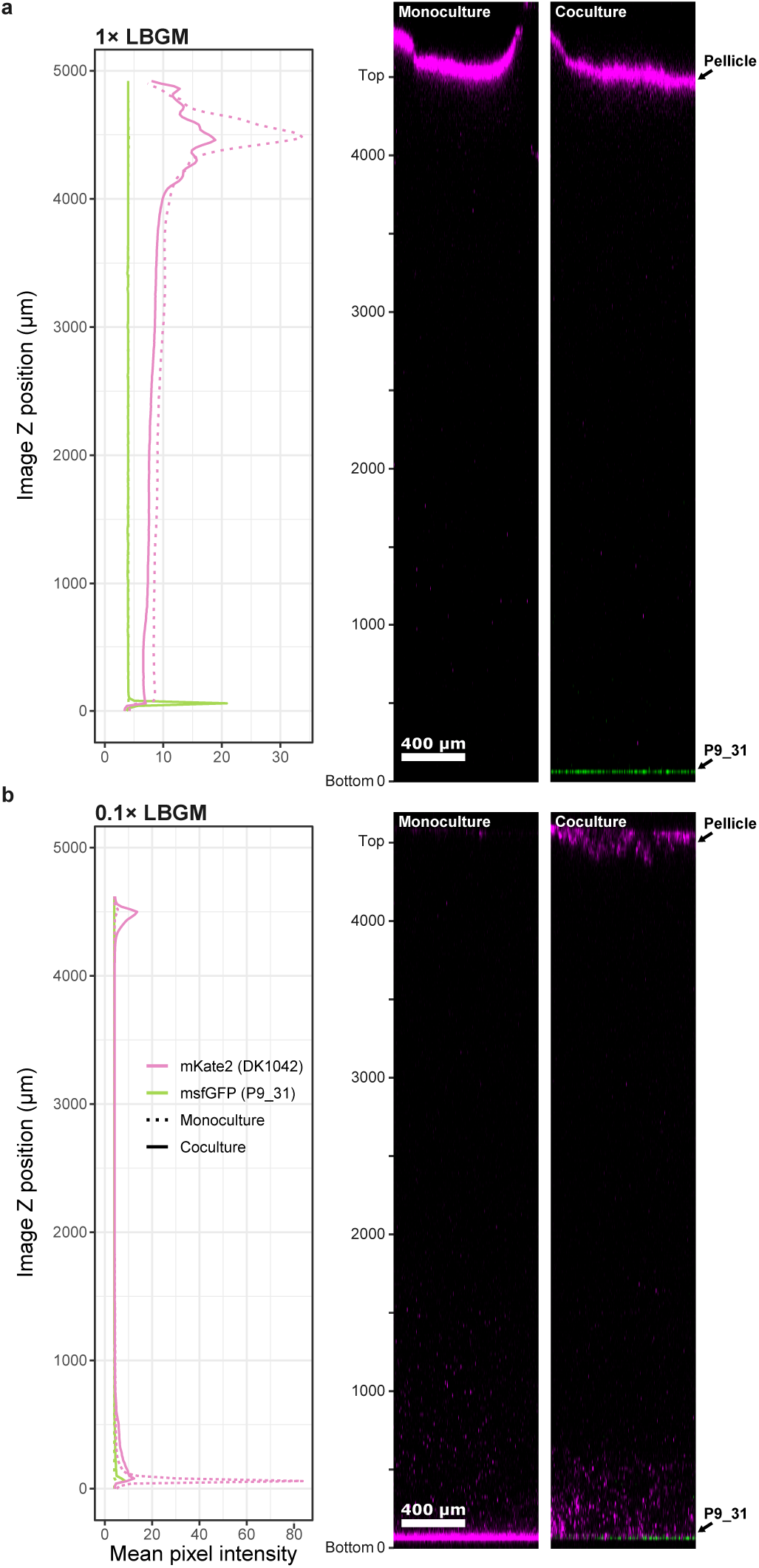
Non-inhibiting *Pseudomonas* is not present in the pellicle. Coculture of DK1042 (magenta) with non-inhibiting isolate P9_31 (green) in 1× LBGM (**a**) and 0.1× LBGM (**b**). Confocal laser scanning microscopy was employed to acquire Z-slices every 20 µm spanning the entire depth of a well. Plots show the mean pixel intensity of each strain over the depth of the well. Pixel intensities cannot be compared between 1× LBGM and 0.1× LBGM.

### DAPG is a biomarker for *B. subtilis* inhibition

Four of five colony-inhibiting isolates were found to contain the conserved *phl* BGC operon known to produce DAPG ^33^. From 61 cocultures on solid 0.1× LBGM, 11 isolates resulted in markedly reduced DK1042 colony growth (Fig. 6a). A PCR screen of the main biosynthetic gene for DAPG, *phlD*, revealed that 10 of 11 colony-inhibiting isolates carried *phlD* (Fig. 6b). By contrast, 0 of 7 non-inhibiting strains tested positive for *phlD*. To determine if DAPG or any intermediate products from the PhlA−D BGC operon, we cocultured *P. protegens* DTU9.1 and mutants lacking *phlACB* (DAPG^-^) or *phlACBD* (phloroglucinol^-^, monoacetylated phloroglucinol^-^, DAPG^-^, chlorinated phloroglucinol^-^ and pyoluteorin^-^) with DK1042 in liquid 1× LBGM and on solid 0.1× LBGM (Fig. 6c). Disrupting the production of DAPG in DTU9.1 markedly improved DK1042 fitness in colonies but not in pellicles. Thus, DAPG is a strong inhibitor of *B. subtilis* growth in colonies and pellicles, but not the only inhibiting factor.

**Fig. 6.**
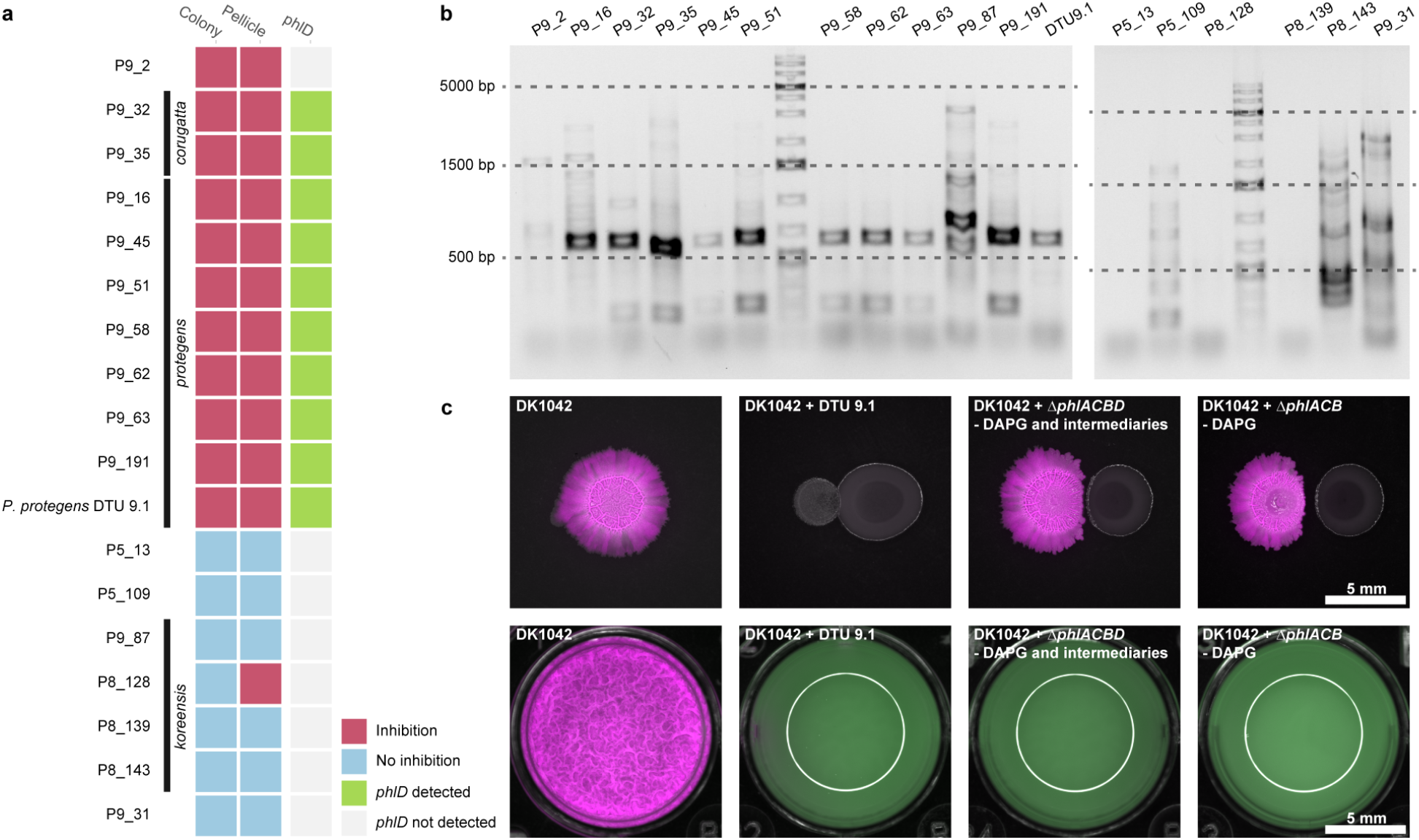
2,4-Diacetylphloroglucinol (DAPG) is a major (but not exclusive) antagonistic metabolite. **a:** Inhibition potential of a selection of isolates and the presence/absence of *phlD* as determined by PCR (**b**). *phlD* was amplified with primers yielding a product of ∼630 bp. Ladder is GeneRuler 1 kb Plus DNA Ladder (Thermo Fisher Scientific). **c:** Cocultures of DK1042 and *P. protegens* DTU 9.1 and *phl*-mutant derivatives of DTU 9.1. Δ*phlACB* cannot produce DAPG. Δ*phlACBD* additionally cannot produce phloroglucinol, monoacetylated phloroglucinol, chlorinated phloroglucinol (PG-Cl/PG-Cl_2_) or pyoluteorin.

## Discussion

In this study, we propose a generalisation of *Bacillus subtilis*-*Pseudomonas* interactions based on *Pseudomonas* taxonomy. Our results suggest that fluorescent pseudomonads are very likely to inhibit the *B. subtilis* pellicle, particularly in 0.1× LBGM. Many studies have demonstrated how nutrient sources determine the outcome of pairwise interactions^14,26,34,35^. Most of our pairwise interactions were negative, fitting the current paradigm. Large-scale pairwise interaction studies have demonstrated that most pairwise interactions are negative, and that the direction of interaction depends on carbon source complexity^26,36,37^. It is interesting, however, that such a large proportion of our interactions proved to be negative or neutral. Granted, our setup only shows the interaction outcome of one of the two participants, hence we cannot differentiate between mutualism (+/+) and parasitism (+/-). Even so, Kehe et al. (2021) reported that for 23% of their >180,000 pairwise interactions at least one participant benefitted from coculturing^26^, while we report only 2.8% positive interactions, making interactions between *B. subtilis* and fluorescent pseudomonads less likely to be positive compared with average interactions among culturable microorganisms.

This could be due to our decision to coculture under pellicle-inducing conditions. DK1042 proved less durable in liquid broth compared with solid agar. It is likely that both the pseudomonads and DK1042 have different metabolic profiles under the two conditions, and that antagonistic molecules are produced in one setting and not the other. Alternatively, the concentration gradients could be more homogeneous under liquid conditions; for example, DK1042 may be unable to reach adequate cell density for biofilm activation before being growth-inhibited. It is likely that a similar screen performed on solid agar media would result in fewer examples of *Pseudomonas-*mediated antagonism.

Predicting interactions from phylogeny is arguably a cornerstone of ecology. Charles Darwin proposed what would become the competition-relatedness hypothesis, stating that closely related species are more likely to compete due to niche overlap^38^. Indeed, relatedness has consistently correlated with competitiveness, though some find that there is a competitive “peak” at intermediate relatedness^39,40^. Bacilli and pseudomonads are phylogenetically distant, belonging to distinct phyla, though metabolically similar enough that syntrophy between isolates has been reported^14^. As such, the competition-relatedness hypothesis states that members of these genera should generally compete. Additionally, these are organisms with large genomes and the potential to produce several antimicrobial compounds. Such organisms cluster into highly competitive communities in metabolic simulations^41^. Additionally, members of *Bacillus* and *Pseudomonas* are frequently co-isolated, suggesting frequent encounters and possible co-evolution. Interactions between *Streptomyces* soil isolates have argued that local evolution is a stronger contributor to interaction outcomes than phylogeny^42,43^. Interactions from a single grain of soil were dramatically different even across isolates with almost identical 16S rDNA sequences, and comparisons between three soil sites found interaction network distributions to differ significantly, again irrespective of phylogenetic distance. In a recent preprint, Pomerleau *et al*. (2023) demonstrated how adaptive laboratory co-evolution of *B. subtilis* with fluorescent pseudomonads increases *B. subtilis* competitive potential^44^. Thus, it is plausible that competitive interactions between bacilli and pseudomonads stem from co-evolution. Predicting interactions from taxonomy requires a certain degree of conservation within taxonomic units. Indeed, with *Pseudomonas* genomes, accessory genes have been found to evolve with core genes, suggesting little horizontal gene transfer, and thus conservation across *Pseudomonas* phylogeny^45,46^. Recently, phylogenetic distance was demonstrated to correlate positively with predictive ability in pairwise interactions, and interactions are conserved within a taxonomic unit but vary between units^34^. We propose that *Pseudomonas* constitutes a taxonomic unit with high phylogenetic predictability, at least in interactions with *Bacillus*. This predictability provides potential for future bioformulations. Both *P. trivialis* and *P. lini* have been implicated in biocontrol and biostimulation^47–49^, and their compatibility with *B. subtilis* therefore implicates them as candidates for beneficial *Bacillus*-*Pseudomonas* consortia. Future studies should investigate how mixed cultures function in the rhizosphere and in more diverse communities.

Positive interactions between *Bacillus* and *Pseudomonas* have been reported before, but never with a fluorescent isolate. Sun et al. (2021) molecularly characterised a mutualistic relationship between *Bacillus velezensis* and *Pseudomonas stutzeri* (now *Stutzerimonas degradans*) ^14^, and described how *B. velezensis* arrives first and subsequently recruits *Pseudomonas* spp. This interaction gave rise to a mixed biofilm of homogeneously distributed *B. velezensis* and *S. degradans*, which we did not observe. Rather, we observed spatial segregation to the air-liquid interface (DK1042) and the liquid-surface interface (P9_31), though still with enhanced pellicle density in 0.1× LBGM. One could speculate that P9_31 secretes a metabolite that can be distributed throughout the medium and that influences *Bacillus* pellicle formation.

We found several antagonistic species, and, like others before us ^50,51^, experimentally demonstrated the influence of DAPG produced by *Pseudomonas* as a key antagonistic metabolite. However, species without the biosynthetic potential to produce DAPG were also characterised as antagonistic, herein members of the *P. putida* group (*P. entomophila* and *P. capeferrum*) and the *P. fluorescens* subgroup. Previous studies demonstrated how cyclic lipopeptide-producing pseudomonads can inhibit *Bacillus* spp. ^52^, and given their potential to produce one or more cyclic lipopeptides, the isolates presented here likely share a similar antagonistic property mediated by bioactive specialised metabolites.

Understanding cocultures of *Bacillus* and *Pseudomonas* is pertinent to their applicability in biotechnology. Especially within agriculture, plant growth-promoting rhizobacteria are being investigated as alternatives to traditional fertilisers and pesticides. Both *B. subtilis* and many fluorescent pseudomonads are characterised as plant growth-promoting, and several products based on members of either genus are currently available ^53,54^. Interestingly, several studies report having successfully combined bacilli and pseudomonads and achieved some form of synergy ^4,55^. Our results suggest that this synergy likely does not stem from increased *Bacillus* growth. This does not contradict the aforementioned studies, which do not report on synergistic growth of members, but on outcomes of other parameters (often plant growth with or without stress). One study even reported biocontrol synergy from a mixture of *B. subtilis* and *P. protegens* ^56^, which suggests either that continued growth of both participants is not required for synergy, or that higher-order interactions abolish DAPG-mediated antagonism of *B. subtilis*.

Both species may not be required to be present at the same time. If the hypothesis proposed by Sun and colleagues that *Bacillus* recruits *Pseudomonas* to the rhizosphere is correct, it is possible that *Pseudomonas* is the effector of biocontrol, eradicating *Bacillus* in the process. However, the fact that many studies have isolated members of both genera from the rhizosphere provides evidence supporting co-existence of the two. Our pairwise interactions then suggest that other factors underpin *Bacillus* and *Pseudomonas* compatibility.

Although future studies are needed to probe the entire breadth of *Pseudomonas* phylogeny and how it enables interaction predictions, our work demonstrates the relevance in doing so.

## Methods

### Culturing and genetic modification

A library of 720 soil isolates was acquired from soil samples taken from pristine grassland in Dyrehaven Lyngby, Denmark, by plating solubilised soil on King’s B agar (KB; 20 g/L peptone, 1% v/v glycerol, 8.1 mmol/L K_2_HPO_4_, 6.08 mmol/L MgSO_4_·7H_2_O) supplemented with 40 µg/mL ampicillin, 13 µg/mL chloramphenicol and 100 µg/mL cycloheximide. Plates were incubated at 30°C for 5 days, and colonies were assessed for fluorescence and placed into lysogeny broth (LB; Lennox, Carl Roth, Karlsruhe, Germany) in 96-well microtiter plates. We labelled the isolation sites as P5 (n = 237 isolates), P8 (n = 279) and P9 (n = 224), where P5 and P9 came from short grass, while P8 came from long grass. DK1042 and soil isolates were routinely cultured in tryptone soy broth (CASO broth; Sigma-Aldrich, Darmstadt, Germany), LB, LB supplemented with 1% glycerol and 0.1 mM MnCl_2_ (1× LBGM), and a 10× dilution of LBGM (0.1× LBGM) at 30°C. Solid media was supplemented with 1.5% (w/v) agar. Antibiotics were added as appropriate in the following final concentrations: gentamycin (Gm) 50 μg/mL, ampicillin (Amp) 100 μg/mL, chloramphenicol (Cm) 10 μg/mL, nalidixic acid (NalAc) 20 µg/mL.

*Pseudomonas* soil isolates were fluorescently tagged by inserting constitutively expressed msfGFP into the attTn7 site as described previously ^57^.

### High-content screening

Precultures of isolates and *B. subtilis* DK1042 Phy-mKate2 were mixed in equal volumes in 96-well imaging microtiter plates (PerkinElmer, Waltham, MA, USA) in TSB or 0.1× LBGM (Fig. 1a). Isolates were adjusted to a final dilution of 100-fold and DK1042 to a final OD_600_ = 0.01. One column contained DK1042 monocultures and one column non-inoculated medium (blank control). Pellicles were incubated at 30°C for 24 h, before removing the supernatant underneath the pellicle and scanning the plate in an Opera QEHS high-content screening microscope (PerkinElmer) equipped with a UAPO20×W3/340 objective with NA = 0.7.

Focal height was adjusted using monoculture samples, and each plate was scanned by imaging four random locations per well, acquiring 21 Z-slices in increments of 2.0 µm from the bottom. Samples were excited with a 561 nm laser collecting emission light through a 690/70 nm filter for mKate2 fluorescence. Laser power was set to 100 µW and samples were excited for 800 ms.

Opera flex-files were imported into FIJI (2.1.0/1.53f51) ^58^ using BioFormats and segmented by applying a 5×5 mean convolution kernel to remove noise, a 3D median filter with radius Z = 2.0 µm to remove single cells or objects only present in one slice, and applying the built-in ImageJ Remove Background function with a rolling ball size of 100 px. Images were then thresholded using the MaxEnthropy algorithm ^59^ based on the pixel intensities in the entire volume (Fig. S1).

Biovolume was calculated using BiofilmQ ^60^ by importing the segmented images and sectioning the segmentation into 8 µm cubes. Objects smaller than 8 µm^3^ were filtered out and the total biovolume from each image stack was determined.

Subsequent analysis was carried out in Rstudio (2022.02.3-b492) ^61^ within R (4.1.1) ^62^ with the Tidyverse framework (1.3.1) ^63^. Log_2_(FoldChange) was calculated between the coculture biovolume of each isolate and the monoculture biovolume in the corresponding plate. Assuming that each participant in a biofilm is theoretically able to occupy 1/k of the space (where k is the number of participants), we divided the biovolume from the monoculture by k to take this lack of space into account.

Isolates were divided into three categories (neutral, negative, or positive) based on the median and interquartile range of three biological replicates (Fig. 2a).

### *rpoD*-targeted amplicon sequencing

To determine the taxonomic composition of the screen output, we employed a targeted amplicon sequencing approach using the species-specific gene *rpoD* as previously demonstrated ^64^.

Precultures of isolates were adjusted to OD = 0.5 and pooled by equal volume according to their category. Genomic DNA was extracted twice from 100 µL of each pool using a Bacterial & Yeast Genomic DNA Purification Kit (EURx, Gdańsk, PL) following the manufacturer’s instructions, yielding 1000−1500 ng of DNA in 50 µL. DNA was also extracted from a pool of all 720 isolates and as a negative control, nuclease-free H_2_O was used in place of a bacterial pool. From each DNA extraction, 10 ng was used as template in two PCR amplification experiments using TEMPase hotstart polymerase (Ampliqon, Odense, DK) and barcoded primer pairs (Table S1), resulting in two PCR mixtures from each of two genomic DNA extractions. A final 25 µL PCR experiment consisted of TEMPase mastermix (1×), forward primer (320 nmol/L), reverse primer (320 nmol/L), MgCl_2_ (1.75 mmol/L), template gDNA and nuclease-free H_2_O. The reaction was initiated with 15 min at 95°C to denature the DNA and activate the TEMPase polymerase, followed by 35 cycles of 30 s denaturation (95°C), 30 s annealing (53°C), 30 s elongation (70°C), and a final 5 min elongation step at 70°C. Read length was assessed by DNA agarose gel electrophoresis (1% w/v) and amplicons were purified from the PCR mix using a NucleoSpin Gel and PCR Clean-up kit (Macherey-Nagel, Düren, DE) following manufacturer’s instructions.

DNA purities and concentrations were assessed using a Denovix spectrophotometer (DeNovix, Wilmington, DE, USA) and a Qubit 2.0 fluorimeter (Thermo Fisher Scientific, Waltham, MA, USA) and a Qubit High Sensitivity kit, respectively. Optical density ratios (260/280 and 260/230) measured 2.0 ± 0.20 and concentrations ranged from 20 to 68 ng/µL. Amplicon DNA (360 ng) from each sample was pooled to a total of 10 µg DNA and sent to Seqomics Biotechnology Ltd. for sequencing on a MiSeq platform (Illumina, San Diego, CA, USA) using a MiSeq Reagent Kit v3 (600-cycle).

The resulting reads were quality-checked and demultiplexed using CutAdapt V4.0 ^65^. FastP V0.23.2 ^66^ was used with default settings for quality filtering. Mapping and post-mapping filtering was performed using the bowtier.sh script from Lauritsen *et al*. 2021 ^64^. In brief, paired reads were aligned with bowtie2 to a custom database of 160 *rpoD* genes. The resulting SAM-file was then filtered for only concordant pairs mapped with a quality >10 using samtools. In Rstudio, a permutational multivariate ANOVA was performed to test for between-sample clustering. Amplicon reads were normalized with DESeq2 ^67^ before calculating relative abundances. Differential abundance analysis was carried out using ANCOM-BC ^68^ with uncorrected read counts and multiple testing was corrected with the false discovery rate (FDR) method ^69^.

In addition, Sanger sequencing of the *rpoD* gene was used for individual routine taxonomic identification of isolates with primers PsEG30F and PsEG790R ^64^.

### Pairwise interactions

On agar, DK1042 *amyE*::P_hyperspank_-*mKate2 sacA*::P_*eps*_-*gfp* and a candidate isolate were spotted 5.0 mm apart on agar surfaces using 2 µL of culture at an OD_600_ of 1.0. Prior to spotting, plates were dried for 30 min in a lateral flow hood then incubated at 30°C for 72 h.

In broth, the DK1042 *amyE*::P_hyperspank_-*mKate2 sacA*::P_*eps*_-*gfp* strain and a candidate isolate were mixed 1:1 volumetrically in 1 mL liquid LBGM in 24-well microtiter plates at a final OD_600_ of 0.01 and incubated for 24 h at 30°C.

Example *Pseudomonas* strains were *P. lini* 1.6, *P. poae* DSM 14936 ^70^, *Pseudomonas kermanshahensis* F8 (previously *P. capeferrum*) ^71^ and *P. protegens* DTU 9.1 ^71^.

### Stereomicroscopy

Colonies and pellicles were imaged with a Carl Zeiss Axio Zoom.V16 stereomicroscope (Carl Zeiss, Oberkochen, Baden-Württemberg, Germany) equipped with a CL 9000 LED light source (Carl Zeiss) and an AxioCam 503 monochromatic camera (Carl Zeiss). The stereoscope was equipped with a PlanApo Z 0.5x/0.125 FWD 114 mm, and the filter sets 38 HE eGFP (ex: 470/40, em: 525/50) and 63 HE mRFP (ex: 572/25, em: 629/62). Exposure time was optimised for contrast but kept constant under identical conditions (i.e. media types or biofilm type).

Image processing and analysis was performed in FIJI. Contrast in fluorescence channels was adjusted identically on a linear scale to allow for visual comparisons between images with equal exposure time. Reporter fluorescence intensity was measured by segmenting the colony of interest with the Triangle thresholding algorithm ^72^ based on mKate2 signal and measuring relative eGFP intensity per area within the resulting region of interest outlining the entire colony. For pellicles, circles were manually fitted to include only the well. Standard deviation of the pixel intensity was measured and reported as a proxy for pellicle wrinkles.

### Confocal microscopy

DK1042 was cultured with isolate P9_31 in 1× LBGM or 0.1× LBGM in 24-well imaging microtiter plates (PerkinElmer). Wells were imaged on a Leica SP8 confocal microscope (Leica Microsystems, Wetzlar, Germany) equipped with an HC PL Fluotar 10x/0.30 air objective and lasers exciting at 488 nm and 552 nm. Photomultiplier tubes were adjusted to acquire photons at wavelengths of 493−581nm (msfGFP) and 586−779nm, and gain was adjusted for optimal contrast, but kept constant for each media type. Images were acquired with 8 bits and size 512px × 512px × 247px (XYZ – voxel size: 1.78 µm ×1.78 µm × 20.00 µm) averaging over four lines.

### Genome mining

Whole-genome sequencing was performed as described previously ^57^. Assembled genomes were annotated with Bakta (V1.6.1) ^73^ and a BGC presence/absence matrix was created with antiSMASH (V7.0.0) ^74^ using its relaxed mode performing KnownClusterBlast, ClusterBlast, SubClusterBlast and MIBiG cluster comparison, and ActiveSiteFinder, RREFinder and TFBS analysis. Results were manually curated using the *Pseudomonas* Genome Database ^75^ as reference. PCR screening for *phlD* was performed with primers B2BF (5’-ACCCACCGCAGCATCGTTTATGAGC) and BPR4 (5’-CCGCCGGTATGGAAGATGAAAAAGTC), yielding a 630 bp product.

## Data availability

Analysis scripts and processed data have been deposited at Github (https://github.com/marklyng/screen_repository). *rpoD* amplicon sequencing data have been deposited at the Sequence Read Archive under BioProject ID PRJNA985909.

## Acknowledgements

This project was funded by a DTU Alliance Strategic Partnership PhD fellowship, by the Danish National Research Foundation (DNRF137) for the Center for Microbial Secondary Metabolites, and the Novo Nordisk Foundation within the INTERACT project of the Collaborative Crop Resiliency Program (NNF19SA0059360), and the “Imaging microbial language in bio-control (IMLiB)” infrastructure grant (NNF19OC0055625).

