## Supplementary material for "Taxonomy of *Pseudomonas* spp determines interactions with *Bacillus subtilis*": Fig S1 to S3 and Table S1

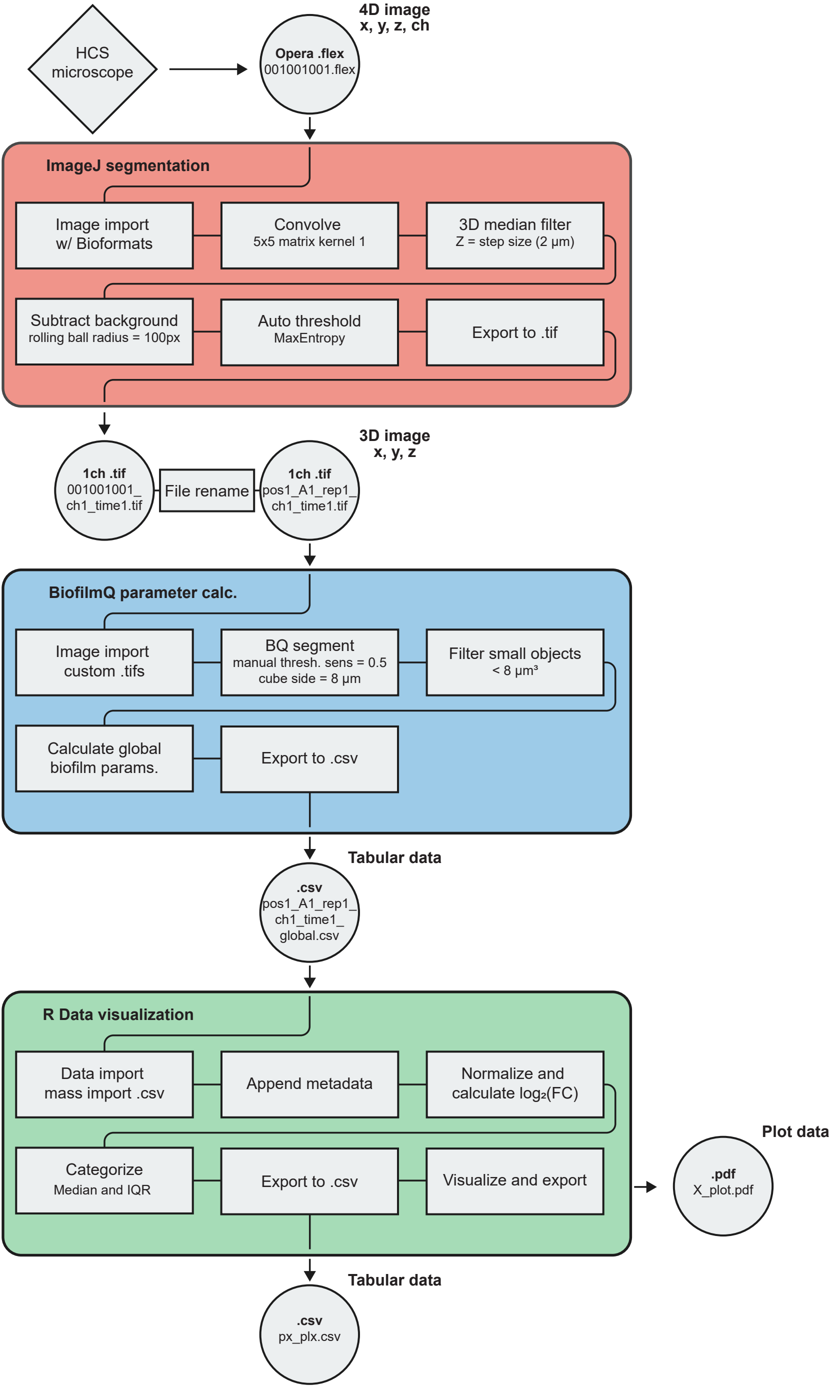

HCS microscope
4D image x, y, z, ch
Opera .flex
001001001.flex
ImageJ segmentation
Image import w/ Bioformats
Convolve
5x5 matrix kernel 1
3D median filter
Z = step size (2 µm)
Subtract background
rolling ball radius = 100px
Auto threshold
MaxEntropy
Export to .tif
1ch .tif
001001001_ch1_time1.tif
File rename
1ch .tif
pos1_A1_rep1_ch1_time1.tif
3D image x, y, z
BiofilmQ parameter calc.
Image import custom .tifs
BQ segment
manual thresh. sens = 0.5
cube side = 8 µm
Filter small objects
< 8 µm³
Calculate global biofilm params.
Export to .csv
Tabular data
.csv
pos1_A1_rep1_ch1_time1_global.csv
R Data visualization
Data import mass import .csv
Append metadata
Normalize and calculate log₂(FC)
Categorize
Median and IQR
Export to .csv
Visualize and export
Plot data
.pdf
X_plot.pdf
Tabular data
.csv
px_plx.csv

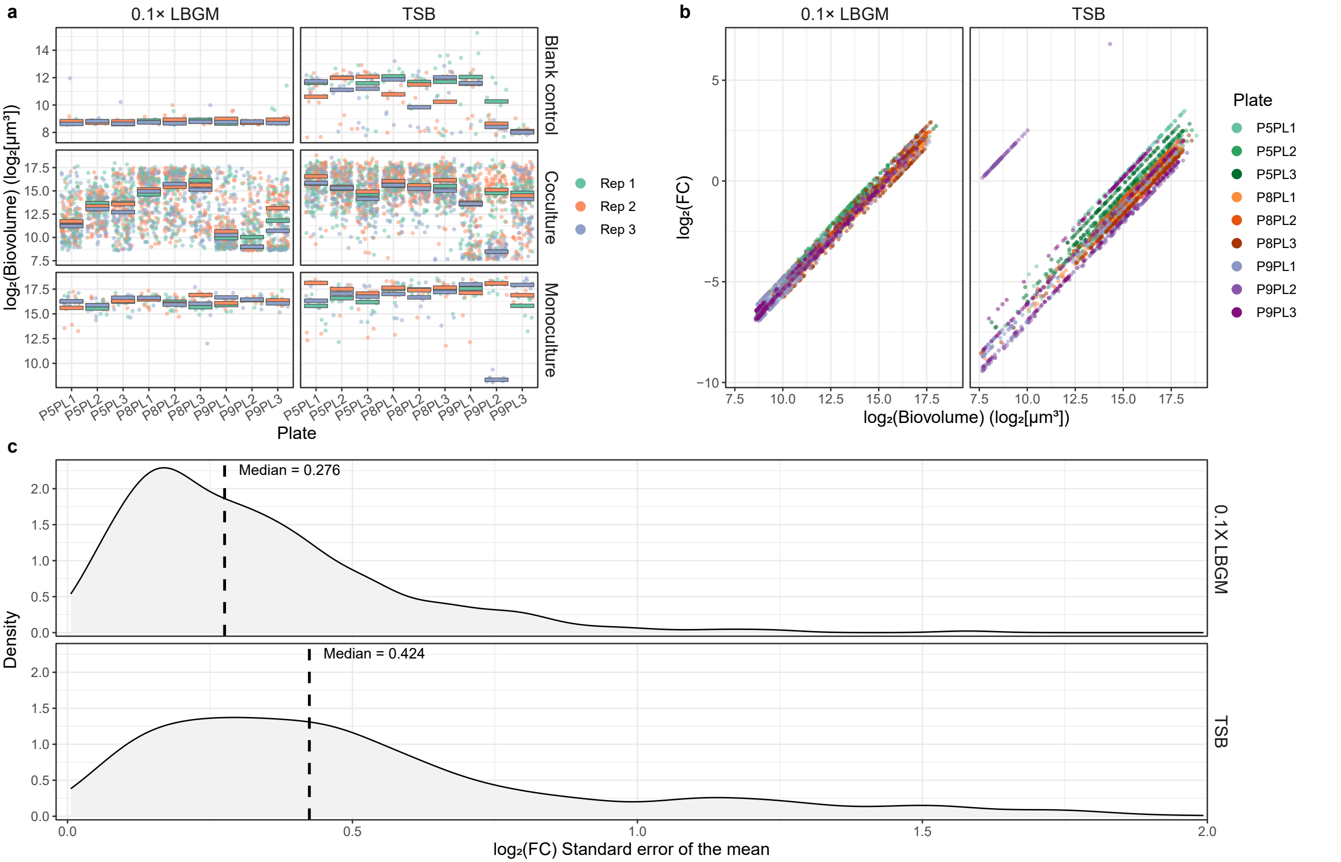
a
0.1× LBGM
TSB
Blank control
Coculture
Monoculture
log₂(Biovolume) (log₂[μm³])
Plate
P5PL1
P5PL2
P5PL3
P8PL1
P8PL2
P8PL3
P9PL1
P9PL2
P9PL3
Rep 1
Rep 2
Rep 3
b
0.1× LBGM
TSB
log₂(FC)
log₂(Biovolume) (log₂[μm³])
Plate
P5PL1
P5PL2
P5PL3
P8PL1
P8PL2
P8PL3
P9PL1
P9PL2
P9PL3
c
Median = 0.276
Median = 0.424
0.1X LBGM
TSB
Density
log₂(FC) Standard error of the mean

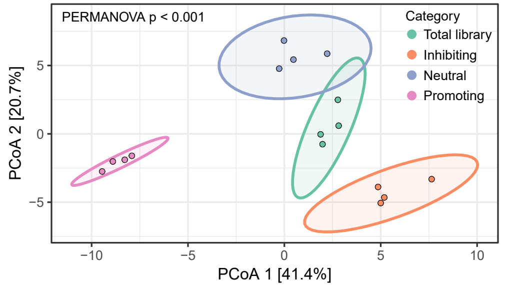
PERMANOVA p < 0.001
Category
Total library
Inhibiting
Neutral
Promoting
PCoA 2 [20.7%]
PCoA 1 [41.4%]
5
0
−5
−10
−5
0
5
10

**Table S1 Strains and oligonucleotides used in the study.**

**Strain**

| ***Bacillus subtilis*** | **Genotype** | **Ref** |
|---|---|---|
| *B. subtilis* DK1042 | 3610 *comI*$^{Q12I}$ | [1] |
| TB501.1 | 3610 *comI*$^{Q12I}$ *amyE*::$P_{hyperspank}$-mKate2-*Spec*$^{R}$ | [2] |
| TB864 | 3610 *comI*$^{Q12I}$ *amyE*::$P_{hyperspank}$-mKate2-*cat*, $P_{eps}$-eGFP-*Km*$^{R}$ | [2] |

| ***Pseudomonas*** | **Genotype** | **Genome Accession** | **Ref** |
|---|---|---|---|
| *Pseudomonas* sp. P5_109 | WT isolate | CP125380 | [3] |
| *Pseudomonas* sp. P5_152 | WT isolate | JASFAH000000000 | [3] |
| *P. zeae* P8_72 | WT isolate | JASFAG000000000 | [3] |
| *Pseudomonas* sp. P8_139 | WT isolate | CP125379 | [3] |
| *Pseudomonas* sp. P8_229 | WT isolate | CP125378 | [3] |
| *Pseudomonas* sp. P8_241 | WT isolate | CP125377 | [3] |
| *Pseudomonas* sp. P8_250 | WT isolate | JASFAF000000000 | [3] |
| *Pseudomonas* sp. P9_31 | WT isolate | CP125375 | [3] |
| *Pseudomonas* sp. P9_2 | WT isolate | CP125376 | [3] |
| *Pseudomonas* sp. P9_32 | WT isolate | CP125374 | [3] |
| *Pseudomonas* sp. P9_35 | WT isolate | CP125373 | [3] |
| *P. germanicum* P9_87 | WT isolate | CP125372 | [3] |
| *P. protegens* P9_191 | WT isolate | JASFAE000000000 | [3] |
| *P. lini* 1.6 | WT isolate | | Unpublished |
| *P. poae* DSM 14936 | Type strain | LT629706 | [4] |
| *P. kermanshensis* F8 | WT isolate | CP099575 | [5] |
| *P. protegens* DTU9.1 | WT isolate | CP024025 | [5] |

| ***Escherichia coli*** | **Genotype** | **Ref** |
|---|---|---|
| *E. coli* CC118 | CC118 λpir/pBG42 | [6] |
| *E. coli* HB101 | HB101 /pRK600 | [6] |
| *E. coli* CC118 | CC118 λpir/pTNS2 | [6] |

| **Oligo ID** | **Description** | **Sequence (5' – 3')** |
|---|---|---|
| PsEG30F-BC13 | Positive 1.1 – Fw | ATTGCTGAATYGAAATCGCCAARCG |
| PsEG30F-BC14 | Positive 1.2 – Fw | TGAGTTCTATYGAAATCGCCAARCG |
| PsEG30F-BC15 | Positive 2.1 – Fw | GGCTATTTATYGAAATCGCCAARCG |
| PsEG30F-BC16 | Positive 2.2 – Fw | CAAGAGATATYGAAATCGCCAARCG |
| PsEG30F-BC17 | Negative 1.1 – Fw | GGAATACAATYGAAATCGCCAARCG |
| PsEG30F-BC18 | Negative 1.2 – Fw | AAGGCAATATYGAAATCGCCAARCG |
| PsEG30F-BC19 | Negative 2.1 – Fw | ACAAAACGATYGAAATCGCCAARCG |
| PsEG30F-BC20 | Negative 2.2 – Fw | TTGAGTGAATYGAAATCGCCAARCG |
| PsEG30F-BC21 | Neutral 1.1 – Fw | GCTTCTGAATYGAAATCGCCAARCG |
| PsEG30F-BC22 | Neutral 1.2 – Fw | GGCAAGATATYGAAATCGCCAARCG |
| PsEG30F-BC23 | Neutral 2.1 – Fw | GTGCTTTCATYGAAATCGCCAARCG |
| PsEG30F-BC24 | Neutral 2.2 – Fw | ACACACTGATYGAAATCGCCAARCG |
| PsEG30F-BC25 | Total 1.1 – Fw | CGATTCTGATYGAAATCGCCAARCG |
| PsEG30F-BC26 | Total 1.2 – Fw | GCAGAGTTATYGAAATCGCCAARCG |
| PsEG30F-BC27 | Total 2.1 – Fw | CGTCCTATATYGAAATCGCCAARCG |
| PsEG30F-BC28 | Total 2.2 – Fw | GCTTGGTTATYGAAATCGCCAARCG |
| PsEG30F-BC29 | Empty control 1 – Fw | ACAGGCTTATYGAAATCGCCAARCG |
| PsEG30F-BC30 | Empty control 1 – Fw | TGACGCTTATYGAAATCGCCAARCG |
| PsEG790R-BC13 | Positive 1.1 – Rv | ATTGCTGACGGTTGATKTCCTTGA |
| PsEG790R-BC14 | Positive 1.2 – Rv | TGAGTTCTCGGTTGATKTCCTTGA |
| PsEG790R-BC15 | Positive 2.1 – Rv | GGCTATTTCGGTTGATKTCCTTGA |
| PsEG790R-BC16 | Positive 2.2 – Rv | CAAGAGATCGGTTGATKTCCTTGA |
| PsEG790R-BC17 | Negative 1.1 – Rv | GGAATACACGGTTGATKTCCTTGA |
| PsEG790R-BC18 | Negative 1.2 – Rv | AAGGCAATCGGTTGATKTCCTTGA |
| PsEG790R-BC19 | Negative 2.1 – Rv | ACAAAACGCGGTTGATKTCCTTGA |

| | | |
|---|---|---|
| PsEG790R-BC20 | Negative 2.2 – Rv | TTGAGTGACGGTTGATKTCCTTGA |
| PsEG790R-BC21 | Neutral 1.1 – Rv | GCTTCTGACGGTTGATKTCCTTGA |
| PsEG790R-BC22 | Neutral 1.2 – Rv | GGCAAGATCGGTTGATKTCCTTGA |
| PsEG790R-BC23 | Neutral 2.1 – Rv | GTGCTTTCCGGTTGATKTCCTTGA |
| PsEG790R-BC24 | Neutral 2.2 – Rv | ACACACTGCGGTTGATKTCCTTGA |
| PsEG790R-BC25 | Total 1.1 – Rv | CGATTCTGCGGTTGATKTCCTTGA |
| PsEG790R-BC26 | Total 1.2 – Rv | GCAGAGTTCGGTTGATKTCCTTGA |
| PsEG790R-BC27 | Total 2.1 – Rv | CGTCCTATCGGTTGATKTCCTTGA |
| PsEG790R-BC28 | Total 2.2 – Rv | GCTTGGTTCGGTTGATKTCCTTGA |
| PsEG790R-BC29 | Empty control 1 – Rv | ACAGGCTTCGGTTGATKTCCTTGA |
| PsEG790R-BC30 | Empty control 1 – Rv | TGACGCTTCGGTTGATKTCCTTGA |
| B2BF | *phlD* – Fw | ACCCACCGCAGCATCGTTTATGAGC |
| BPR4 | *phlD* – Rv | CCGCCGGTATGGAAGATGAAAAAGTC |
| PsEG30F | *rpoD* – Fw | ATYGAAATCGCCAARCG |
| PsEG790R | *rpoD* – Rv | CGGTTGATKTCCTTGA |

Underlined: Barcode for amplicon sequencing.
PsEG30F, PsEG790R, and their barcoded derivatives are from [7].
